# Transcriptomic analysis reveals SnTox8-mediated reprogramming of wheat defence signalling

**DOI:** 10.64898/2026.07.02.736222

**Authors:** Asiri A. Padukka Vidanalage, Kristina K. Gagalova, Eiko Furuki, Fiona Kamphuis, Kasia Rybak, Sambasivam Periyannan, Mark Gibberd, Huyen T. T. Phan

## Abstract

*Parastagonospora nodorum* (Berk.) Quaedvlieg, Verkley & Crousis, a necrotrophic fungal pathogen, is the causal agent for septoria nodorum blotch, a major constraint on global wheat production. Pathogen-produced necrotrophic effectors (NEs) that interact with host-sensitivity genes in an inverse gene-for-gene manner, collectively leading to effector-triggered susceptibility (ETS). Here, we investigated the transcriptional responses of two *Triticum aestivum* L. genotypes, Mace and Lancer, following infiltration with a novel NE, SnTox8. A total of 12,679 unique differentially expressed genes in Mace and 149 in Lancer were detected from transcriptomic analysis. In the SnTox8-sensitive cultivar, Mace, numerous defence-related genes were induced, including protein phosphorylation cascades, reactive oxygen species bursts, calcium signalling, phytohormone modulation, and suppression of photosynthesis, consistent with findings from other ETS models, in which necrotrophic fungal pathogens hijack host defence systems to proliferate. The interaction also activated genes involved in signal transduction, metabolism, membrane modification, and molecular transport, reflecting a coordinated host reprogramming that promotes cellular dysfunction and cell death, thereby facilitating necrotrophic pathogenesis. In contrast, Lancer, an SnTox8-insensitive cultivar, exhibited minimal transcriptional changes with no evidence of effector recognition or downstream defence-related activities. Overall, this study exhibited that SnTox8 manipulates kinase-mediated immune signalling and metabolic reprogramming to convert defence activation into host cell death, revealing a mechanistic basis for ETS in wheat. The identified SnTox8-*Snn8*-triggered processes were confirmed through additional transcriptome analysis of Mace mutants. Outcomes from this study establish a foundation for identifying, functionally characterising and validating the corresponding host susceptibility gene *Snn8*.

## INTRODUCTION

Plants, when exposed to biotic stresses, trigger a cascade of molecular, biochemical, and physiological processes that activate diverse defence signalling pathways. This complex network of responses allows plants to adapt and resist infection by coordinating multiple layers of their immune system. Plants have two defence mechanisms: the first line of defence starts with the recognition of pathogen-associated molecular patterns (PAMPS) via pattern recognition receptors (PRRs), which induces intracellular signalling pathways leading to the activation of PAMP-triggered immunity (PTI) (Jones & Dangl, 2006). Pathogens secrete molecules called effectors to counteract PTI and promote infection. In cases where pathogens can suppress PTI, plants have developed a second layer of defence, which is the recognition of pathogen-produced effectors via plant resistance genes, leading to stimulation of effector-triggered immunity (ETI)(Dodds & Rathjen, 2010). Both PTI and ETI play crucial roles in stimulating the plant defence response against the pathogen, and ultimately leading to a hypersensitive cell death response surrounding the pathogen activity (Jones & Dangl, 2006). Biotrophic and hemibiotrophic pathogens fit well into this ‘gene-for-gene’ defence model, except necrotrophic pathogens, which follow the inverse gene-for-gene interaction. These pathogens hijack the plant defence response, triggering plant cell death to survive (Gao et al., 2015).

*Parastagonospora* (syn. *Stagonospora*; *Phaeosphaeria*, *Septoria*) *nodorum* (Berk.) Quaedvlieg, Verkley & Crousis, a necrotrophic fungal pathogen, is the causal agent of septoria nodorum blotch (SNB), one of the major foliar and glume diseases affecting wheat crops worldwide, including Australia (Peters Haugrud et al., 2022), it impacts wheat yield, grain quality, and quantity by reducing photosynthetic capacity and leading to shrivelled seeds. Knowledge of the biological processes involved in wheat’s responses to SNB has expanded significantly in recent years. Genetic and biochemical studies have uncovered significant interactions between wheat and *P. nodorum*, enhancing understanding of how the disease develops and progresses. One of the breakthroughs in SNB research has been the identification of specific necrotrophic effectors (NE) produced by *P. nodorum* and their interactions with corresponding host-sensitivity genes in wheat, triggering plant cell death and leading to disease development (Li et al., 2021). Such SNB research is critical for elucidating the molecular mechanisms underlying SNB susceptibility and for developing effective strategies to improve disease resistance in wheat.

To date, eleven interactions have been discovered between *P. nodorum* effectors (SnTox) and corresponding sensitivity host genes (*Snn*), including SnToxA-*Tsn1* (Faris et al., 2010; Friesen et al., 2009; Liu et al., 2006), SnTox1-*Snn1(Snn1-B1 & Snn1-B2)* (Liu et al., 2004; Seneviratne et al., 2024; Shi et al., 2016), SnTox267-*Snn2* (Friesen et al., 2009; Friesen et al., 2007; Richards et al., 2021; Zhang et al., 2009), SnTox3-*Snn3-B1* (Friesen et al., 2008; Liu et al., 2009), SnTox3-*Snn3-B2* (Zengcui Zhang et al., 2025), SnTox3-*Snn3-D1* (Friesen et al., 2008; Liu et al., 2009), SnTox4-*Snn4* (Abeysekara et al., 2012; Abeysekara et al., 2009), SnTox5-*Snn5* (Friesen et al., 2012; Kariyawasam et al., 2021), SnTox267-*Snn*6 (Gao et al., 2015; Richards et al., 2021), and SnTox267-*Snn7* (Peters Haugrud et al., 2022; Richards et al., 2021). To date, five host susceptibility genes, *Tsn1* (Faris et al., 2010), *Snn1* (Shi et al., 2016), *Snn3-D1* (Zhang et al., 2021), *Snn3-B1*, and *Snn3-B2* (Z. Zhang et al., 2025) have been cloned along with five NE genes from the pathogen, *SnToxA*, *SnTox3*, *SnTox1*, *SnTox5*, and *SnTox267* (Choupannejad et al., 2024; Kariyawasam et al., 2021). In addition to the major NE-*Snn* interactions, previous studies have reported additional QTLs, suggesting unidentified NE-*Snn* interactions (Peters Haugrud et al., 2022). The identification and cloning of these genes and their interactions have significant implications for breeding disease-resistant wheat varieties. Investigating differentially expressed genes (DEGs) through transcriptomic analysis during pathogen infection provides critical insights into the signalling pathways involved in plant immune responses. Numerous studies have used this approach to explore the molecular mechanisms underlying plant defence. In the case of wheat’s interaction with *P. nodorum*, understanding these pathways is crucial for unravelling the mechanisms that govern disease resistance and susceptibility, which helps to design suitable strategies for resistance deployment in wheat breeding programs.

Early works used whole-leaf microarrays to profile wheat responses to necrotrophic effectors. For PtrToxA (ToxA), transcriptional profiling of ToxA-treated leaves of a ToxA-sensitive wheat cultivar was performed using the GeneChip Wheat Genome Array by Pandelova et al (2009). This study has revealed that rapid induction of defence-associated transcripts (up-regulation of pathogenesis-related genes, activation of MAPK and calcium signalling cascades, etc), alongside ToxA-induced cell death, is triggered by impairment of the photosynthetic machinery and accumulation of reactive oxygen species (Pandelova et al., 2009). A similar study by Adhikari et al (2009) using the Affymetrix GeneChip Wheat Genome Array to investigate the ToxA-*Tsn1* interaction revealed genes associated with the phenylpropanoid pathway, lignification, and ROS generation. Furthermore, the accumulation of H_2_O_2_ in the leaves undergoing programmed cell death provided strong evidence that ROS signalling plays a central role in mediating ToxA-induced cell death during pathogen attack (Adhikari et al., 2009; Tan et al., 2010). A study on SnTox3, combining microarray and proteomics, showed a strong defence signature, including pathogenesis-related genes, and the induction of cell death, strong induction of enzymes involved in primary metabolism, and impairment of photosynthetic processes. Moreover, several compounds associated with plant defence were inducted, including the phenylpropanoids, chlorogenic acid and feruloylquinic acid, as well as the cyanogenic glucoside dhurrin in susceptible wheat leaves (Winterberg et al., 2014). Importantly, these findings supported the view that perception of necrotrophic effectors can activate defence pathways normally associated with resistance to biotrophic pathogens, emphasising how *P. nodorum* exploits host immune signalling to promote disease development. Together, these studies illustrated how transcriptomic and proteomic technologies have progressively improved our understanding of the complex molecular responses causing necrotrophic effector-triggered susceptibility in wheat.

Current studies employed high-throughput RNA-seq approaches to capture dynamic transcriptional reprogramming during plant-pathogen interactions with greater sensitivity and accuracy. For example, Xiong et al. (2018) performed a transcriptomic analysis on *Botrytis cinerea*, the causal agent of “Gray Mold” disease in mature strawberry fruit. Their study revealed that DEGs were predominantly associated with pathogen recognition, signal transduction (MAPK cascades and CDPKs), defence responses (including transcription factors, secondary metabolite biosynthesis, and pathogenesis-related), and various transporter-encoding genes (Xiong et al., 2018). Similarly, a transcriptomic profiling study conducted on both resistant and susceptible wheat varieties infected with *Rhizoctonia cerealis*, the causative agent of sheath blight disease, demonstrated that DEGs were enriched in pathways such as secondary metabolite biosynthesis, carbon metabolism, plant hormone signalling, and plant–pathogen interactions. Notably, the phenylpropane biosynthesis pathway was specifically activated in the resistant variety post-infection. Many DEGs belonged to the MYB, AP2, NAC, and WRKY transcription factor families (Geng et al., 2022). Another study focused on soybean cyst nematode (SCN), a root-parasite of soybean, and compared gene expression in resistant and susceptible genotypes across two SCN types. The findings revealed distinct transcriptomic responses, including both commonly and oppositely regulated genes. In the susceptible genotype, the expression levels of shared regulated genes were insufficient to mount an effective defence response. Further analysis indicated that genetic variability, metabolic and hormonal differences, and varied regulation of protein phosphorylation and ubiquitination shaped genotype-specific responses. In contrast, shared responses were driven by conserved immune signalling and SCN-responsive genes, highlighting the complexity and specificity of plant immune strategies (Sultana et al., 2024).

In the present study, the transcriptional responses of two *Triticum aestivum* (wheat) genotypes, sensitive and insensitive to the novel effector SnTox8 secreted by *P. nodorum,* were investigated using mRNA sequencing data. The aims of this study were: (1) to identify main molecular pathways activated during exposure to SnTox8, (2) compare the transcriptional responses of two wheat genotype’s contrasting interactions between *T. aestivum* and *P. nodorum* effector SnTox8; (3) to validate these key molecular pathways using loss-of-function mutants; (4) compare the findings with other NE-receptor interactions in the same pathosystem to evaluate consistency and identify shared or distinct mechanistic features. Findings from this study will provide genetic resources and references for research into the molecular mechanisms of SNB resistance in wheat and its relatives, as well as for breeding SNB-resistant varieties.

## MATERIALS AND METHODS

First, we conducted a preliminary experiment to assess phenotypic differences, both morphological and microscopic, between sensitive and insensitive varieties following infiltration with SnTox8 and the control. Then, we conducted two key RNA-seq experiments:

(1) RNA-seq analysis of both sensitive and insensitive varieties to investigate temporal gene expression responses and signalling pathways, and (2) an RNA-seq experiment using Mace mutants to validate the results of the first RNA-seq experiment at 2 days post-infiltration (dpi) of SnTox8.

### Treatment preparation: SnTox8 and control

SnTox8 was produced in *E. coli* using the pET expression system (Novagen) with the six-histidine tag on its C-terminus (Phan et al., 2018; Tan et al., 2012). Effector production and purification are as described in Furuki et al (submitted). Purified protein was dialysed into filter-sterilised 20 mM sodium phosphate buffer, pH 7.0. The working concentration for plant bioassays was 0.1 mg/ml. Empty vector (pET21a) culture filtrate was used as a negative control. Effectors were snap-frozen and stored at -80 °C.

### Plant material

Sensitive wheat line (Mace) and insensitive line (Lancer) in response to SnTox8, a recently identified *P. nodorum* effector (under-preparation, Furuki et al), were selected for the experiment. The Mace wheat variety, developed by Australian Grain Technologies (AGT) from a cross between Wyalkatchem and Stylet, and was rapidly adopted by Australian growers following its release in 2008 (Moffat et al., 2015). Lancer variety was developed by LongReach Plant Breeders, with the pedigree designation LPB07-0548 (https://www.longreachpb.com.au/products/longreach-lancer, accessed on 2/09/2025).

### Widefield microscopic observations

First leaves of 10-day-old Mace and Lancer seedlings were infiltrated with SnTox8 and a control (empty vector). After infiltration, seedlings were placed in a Conviron with a 12 h photoperiod. Leaf samples (∼3 cm in length) were randomly collected from the infiltrated area at four time points: day 0, day 4, day 7, and day 10 post-infiltration. At each time point, leaf samples were grouped into Petri dishes by wheat cultivar, treatment, time point, and preparation method. Each petri dish, containing 6-8 leaf samples for both sample preparation methods (3-4 replicates for each method), was subsequently processed as follows:

### Fresh leaf sample preparation

Leaf samples were soaked in 96% ethanol for 24 hours, then replaced with fresh 96% ethanol and incubated for an additional 48 hours. On the final day, 3 to 4 samples from each Petri dish were placed on slides, excess ethanol was wiped off, and a few drops of 60% glycerol were added. Coverslips were applied carefully to avoid air bubbles (Nelson et al., 2023). These samples were observed using an Olympus Widefield Microscope.

### Boiled leaf sample preparation

Remaining leaf samples in the 96% ethanol Petri dishes (3-4 samples)) were placed individually into 2-ml centrifuge tubes containing 10% KOH and incubated at 85°C for 2 minutes. Subsequently, the samples were transferred to 2 ml tubes containing 1× phosphate-buffered saline (PBS, pH 7.4) and rinsed three times with PBS. Prepared leaf samples were then placed on slides, excess PBS was wiped, and 60% glycerol was added before applying coverslips (Nelson et al., 2023).

Finally, at each time point, all samples (Mace and Lancer infiltrated with SnTox8 and control) that were treated with both fresh and boiled methods were observed using an upright Olympus widefield microscope (Bright Field mode) at 10x magnification. A contrast setting of 10 ms was used for fresh leaves and 7 ms for boiled leaves.

### RNA-seq study (RNA-seq experiment 1)

#### Growth conditions of planting material

Wheat seeds, both Mace and Lancer, were sown in 30-well trays filled with vermiculite and grown at 21 °C under a 12-hour photoperiod. The experimental design incorporated a completely randomised design with two treatment types - variety (Lancer or Mace) and infiltration (SnTox8 or control), and three replicates.

#### Plant treatment and sample collection

For each wheat variety,12-day-old seedlings were infiltrated with SnTox8 or a solution containing the empty vector (control) using a 1-mL syringe without a needle. The infiltration boundaries on each leaf were marked using a nonphytotoxic marker pen. Leaf tissue samples were collected from the infiltrated areas. Sampling was performed for both genotypes under two treatment conditions, with three biological replicates per condition. The experiment included five time points: 0, 2, 3, 5, and 7 days (Leaf samples were collected at multiple time points following SnTox8 infiltration to capture temporal changes in gene expression associated with effector-triggered responses). This design yielded 60 samples in total. Immediately following collection, all RNA was snap-frozen in liquid nitrogen and stored at -80 °C until extraction.

#### RNA extraction and sequencing

RNA was extracted from leaf samples using the RNeasy Mini Kit (QIAGEN Pty Ltd, PO Box 169, Chadstone Centre, Victoria 3148, Australia). RNA quality was assessed by gel electrophoresis, and concentration was measured using the Qubit™ RNA BR Assay Kit (Invitrogen). All samples showed sufficient RNA quality and concentration for downstream applications, with concentrations ranging from 26 to 158 ng/μl. Residual DNA was removed using the DNA-Free™ Kit (Invitrogen by Thermo Fisher Scientific Australia, 5791 Van Allen Way, Carlsbad, CA 92008, USA). The purified RNA samples were subsequently submitted to the Australian Genome Research Facility (AGRF) in Western Australia for sequencing. An overview of the experimental setup (Experiment 1) is displayed in Table 1.

**Table 1:** Overview of the RNA-seq-based experimental setup for expression analysis.

| No | Sample ID | Timepoint | Cultivar | Treatment | Replicate | No | Sample ID | Timepoint | Cultivar | Treatment | Replicate |
| --- | --- | --- | --- | --- | --- | --- | --- | --- | --- | --- | --- |
| 1 | HPR317 | 0 dpi | Mace | Pet | R1 | 31 | HPR321 | 0 dpi | Lancer | Pet | R1 |
| 2 | HPR318 | 0 dpi | Mace | Pet | R2 | 32 | HPR322 | 0 dpi | Lancer | Pet | R2 |
| 3 | HPR319 | 0 dpi | Mace | Pet | R3 | 33 | HPR323 | 0 dpi | Lancer | Pet | R3 |
| 4 | HPR325 | 0 dpi | Mace | SnTox8 | R1 | 34 | HPR329 | 0 dpi | Lancer | SnTox8 | R1 |
| 5 | HPR326 | 0 dpi | Mace | SnTox8 | R2 | 35 | HPR330 | 0 dpi | Lancer | SnTox8 | R2 |
| 6 | HPR327 | 0 dpi | Mace | SnTox8 | R3 | 36 | HPR331 | 0 dpi | Lancer | SnTox8 | R3 |
| 7 | HPR373 | 2 dpi | Mace | Pet | R1 | 37 | HPR377 | 2 dpi | Lancer | Pet | R1 |
| 8 | HPR374 | 2 dpi | Mace | Pet | R2 | 38 | HPR378 | 2 dpi | Lancer | Pet | R2 |
| 9 | HPR375 | 2 dpi | Mace | Pet | R3 | 39 | HPR379 | 2 dpi | Lancer | Pet | R3 |
| 10 | HPR381 | 2 dpi | Mace | SnTox8 | R1 | 40 | HPR385 | 2 dpi | Lancer | SnTox8 | R1 |
| 11 | HPR382 | 2 dpi | Mace | SnTox8 | R2 | 41 | HPR386 | 2 dpi | Lancer | SnTox8 | R2 |
| 12 | HPR383 | 2 dpi | Mace | SnTox8 | R3 | 42 | HPR387 | 2 dpi | Lancer | SnTox8 | R3 |
| 13 | HPR397 | 3 dpi | Mace | Pet | R1 | 43 | HPR401 | 3 dpi | Lancer | Pet | R1 |
| 14 | HPR398 | 3 dpi | Mace | Pet | R2 | 44 | HPR402 | 3 dpi | Lancer | Pet | R2 |
| 15 | HPR399 | 3 dpi | Mace | Pet | R3 | 45 | HPR403 | 3 dpi | Lancer | Pet | R3 |
| 16 | HPR405 | 3 dpi | Mace | SnTox8 | R1 | 46 | HPR409 | 3 dpi | Lancer | SnTox8 | R1 |
| 17 | HPR406 | 3 dpi | Mace | SnTox8 | R2 | 47 | HPR410 | 3 dpi | Lancer | SnTox8 | R2 |
| 18 | HPR407 | 3 dpi | Mace | SnTox8 | R3 | 48 | HPR411 | 3 dpi | Lancer | SnTox8 | R3 |
| 19 | HPR421 | 5 dpi | Mace | Pet | R1 | 49 | HPR425 | 5 dpi | Lancer | Pet | R1 |
| 20 | HPR422 | 5 dpi | Mace | Pet | R2 | 50 | HPR426 | 5 dpi | Lancer | Pet | R2 |
| 21 | HPR423 | 5 dpi | Mace | Pet | R3 | 51 | HPR427 | 5 dpi | Lancer | Pet | R3 |
| 22 | HPR429 | 5 dpi | Mace | SnTox8 | R1 | 52 | HPR433 | 5 dpi | Lancer | SnTox8 | R1 |
| 23 | HPR430 | 5 dpi | Mace | SnTox8 | R2 | 53 | HPR434 | 5 dpi | Lancer | SnTox8 | R2 |
| 24 | HPR431 | 5 dpi | Mace | SnTox8 | R3 | 54 | HPR435 | 5 dpi | Lancer | SnTox8 | R3 |
| 25 | HPR437 | 7 dpi | Mace | Pet | R1 | 55 | HPR441 | 7 dpi | Lancer | Pet | R1 |
| 26 | HPR438 | 7 dpi | Mace | Pet | R2 | 56 | HPR442 | 7 dpi | Lancer | Pet | R2 |
| 27 | HPR439 | 7 dpi | Mace | Pet | R3 | 57 | HPR443 | 7 dpi | Lancer | Pet | R3 |
| 28 | HPR445 | 7 dpi | Mace | SnTox8 | R1 | 58 | HPR449 | 7 dpi | Lancer | SnTox8 | R1 |
| 29 | HPR446 | 7 dpi | Mace | SnTox8 | R2 | 59 | HPR450 | 7 dpi | Lancer | SnTox8 | R2 |
| 30 | HPR447 | 7 dpi | Mace | SnTox8 | R3 | 60 | HPR451 | 7 dpi | Lancer | SnTox8 | R3 |

#### RNA-seq analysis

A minimum of 50M 150bp-PE short reads from RNA were sequenced for each sample using the Illumina NovaSeq sequencing technology (Illumina, Inc., San Diego, USA). The sequencing reads were evaluated with FastQC v0.11.9 (Simon Andrews et al., 2010) and subsequently trimmed using trimmomatic (Bolger et al., 2014). The quality of the trimmed reads was retested with FastQC (fastqc/0.11.9--hdfd78af_1, Andrews (2010)) and values of >= 30 were used for further analysis. Cleaned reads were mapped to the corresponding reference genomes using the hisat2 v2.2.1 package (Kim et al., 2019): for the sensitive line, the Ensembl assembly PGSBv2.1 (accession: GCA_903994175.1, cultivar Mace, accessed on 01/07/2025), and for the insensitive line, the Ensembl assembly PGSBv2.1 (accession: GCA_903993975.1, cultivar Lancer, accessed on 01/07/2025). The aligned sequencing reads were stored as BAM (Binary Alignment/Map) files and subsequently visualised in Integrative Genomics Viewer (IGV) to inspect sequence variations in candidate genes. Bam files generated from the mapping step were used to produce read-count tables for all samples using the feature Counts function in subread v2.0.3 (Liao et al., 2014).

All samples were subjected to exploratory data analysis to assess the overall structure and variability of the data. Principal Component Analysis (PCA) was performed and visualised through PCA plots to identify potential clustering patterns among treatments and time points, for each genotype.

Differential gene expression analysis was performed using the *DESeq2* package (version 1.49.2; Love et al., 2014) in R (version 4.3.2). To account for the time-series structure of the experiment measurements, a reduced-model approach was employed using the Likelihood Ratio Test (LRT). Because the two wheat genotypes were analysed separately, variety was not included as a factor in the statistical model. Therefore, the full model included treatment, timepoint, and their interaction (∼treatment + timepoint + treatment: timepoint). Timepoints were included to capture dynamic transcriptional responses following SnTox8 infiltration, and the 0-dpi time point served as the baseline for comparison with later time points. Prior to analysis, low-count genes were filtered out by retaining only those with counts of at least 10 in three or more samples. The reduced model excluded the interaction term (∼ treatment + timepoint), and the LRT was used to test for the significance of the interaction effect. Subsequent pairwise Wald tests were performed to extract the differential expression results at specific time points, comparing the interaction between the SnTox8 treatment and control at each time point. Genes were considered significantly differentially expressed (DEGs) if they exhibited a Benjamini-Hochberg adjusted *p*-value < 0.01 and an absolute log2 fold change > 1.

The resulting DEGs were visualised using the *EnhancedVolcano* package, version 1.13.2 (Blighe, 2025) to generate volcano plots for each time point, for intuitive interpretation of statistical significance. Additionally, overlaps among DEGs across different time points were visualised using the *UpSetR* package (Conway et al., 2017), allowing identification of shared and unique expression responses over the time course. The Gene Ontology (GO) enrichment analysis of DEGs was conducted by using the topGO R package, version 2.54.0, with Fisher’s exact statistic in the weight algorithm (Alexa & Rahnenführer, 2009). The analysis was conducted using the *GO.db* annotation package, version 3.18.0. GO term annotations for both genotypes were retrieved using the *biomaRt* in Plant Ensemble database (https://plants.ensembl.org/Triticum_aestivum) (PGSBv2.0). GO terms were considered enriched if the associated adjusted p-value was lower than 0.05.

Heatmaps representing expression patterns of DEGs across samples were generated using the *ComplexHeatmap* package, version 2.18.0. Both genes and samples were clustered using Euclidean distance and the Ward D2 method. For Mace, a DESeq2 dataset was modelled with treatment, timepoint, and their interaction. After a variance stabilising transformation, all significant genes were extracted and clustered with degPatterns from DEGreport, version 1.38.5 (Pantano, 2017), using treatment and timepoint to capture dynamic transcriptional responses to SnTox8.

### Validation by Mutagenesis (RNA-seq experiment 2)

Seeds of the wheat variety Mace were treated with ethyl methane sulfonate (EMS) following the protocol described by Periyannan (Periyannan & Periyannan, 2017). An initial dosage curve analysis indicated that 0.4% (v/v) EMS was the optimal mutagenic concentration, resulting in approximately 50% seed germination and a 50% reduction in plant height. Then, nearly 2000 seeds were pre-soaked in water for 16 h with orbital rotation at 4 ^0^C. Then, the water-drained seeds were stored at 4 ^0^C for a few hours and treated by soaking at room temperature for 12 h in a 0.4 % (v/v) EMS solution. The EMS solutions were aerated by gentle agitation on a shaker during treatment. Treated seeds were added to 100 mM sodium thiosulfate at the same volume and shaken for 15 minutes. After repeating the sodium thiosulfate treatment twice, the seeds were rinsed with water 3 times. Treated seeds were then rinsed in running tap water for 1 h to remove the EMS solution from the surfaces, left to air dry for 1 h, and immediately planted to obtain M_1_ plants. Accordingly, 3800 heads (M_2_ seeds) were harvested separately from the cultivation of 1008 M_1_ plants, and the planting of new generation seeds continued to obtain M_3_ progeny. 10-day-old M_3_ progeny seedlings were subsequently infiltrated with the SnTox8 effector, and plants were scored for disease development. M_3_ plants that exhibited an insensitive response were transplanted and self-pollinated to produce M_4_ seeds, which were planted and re-screened with SnTox8 to confirm their phenotype. Then, RNA was extracted from three SnTox8-insensitive mutants and one SnTox8-sensitive ectopic/sister-mutant samples from these M_4_ progeny, along with wild-type *Mace*. Samples were collected at Day 2 post-infiltration (2 dpi), with 2 biological replicates per time point, for a total of 10 samples. All samples were snap-frozen in liquid nitrogen and stored at -80 °C until RNA extraction. RNA was extracted as described in section 3.4.3. A minimum of 50M 150bp-PE short reads from total mRNA were obtained for each sample, and short reads were evaluated with FastQC and subsequently trimmed as described previously in section 3.4.4. The cleaned reads were mapped to the corresponding reference genome Mace. Finally, Bam files generated during the mapping step were used to produce read-count tables for all samples.

Expression analysis of RNA-seq data from three loss-of-function mutants (insensitive Mace) and one sister mutant (SnTox8 sensitive) was conducted using the DESeq2 package in R as described above. Table 2 displayed the details of the samples, replicates, and time points of experiment 2. Day 2 mutant experiment data were compared with Day 2 RNA-seq data from the first experiment to identify changes in gene expression associated with the SnTox8 response among mutants and wild-type Mace plants.

**Table 2:**
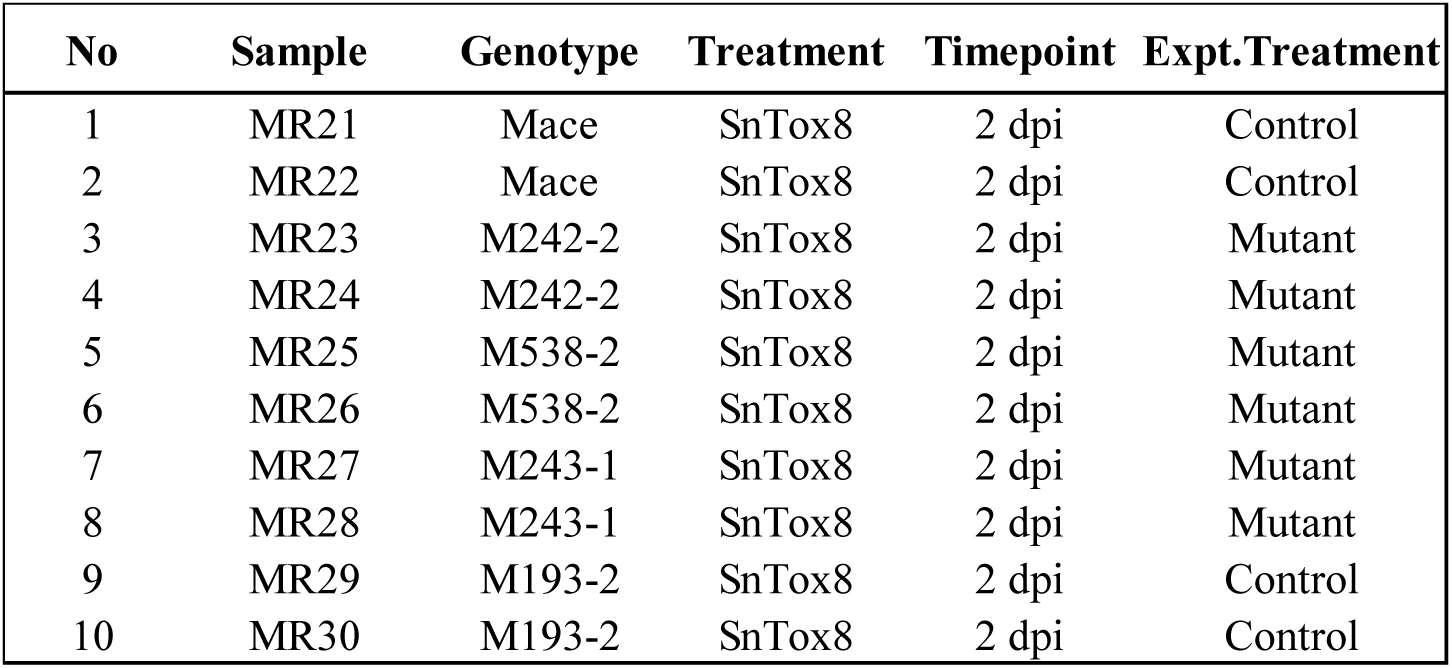
Overview of the mutant-based experimental setup for expression analysis.

| No | Sample | Genotype | Treatment | Timepoint | Expt.Treatment |
| --- | --- | --- | --- | --- | --- |
| 1 | MR21 | Mace | SnTox8 | 2 dpi | Control |
| 2 | MR22 | Mace | SnTox8 | 2 dpi | Control |
| 3 | MR23 | M242-2 | SnTox8 | 2 dpi | Mutant |
| 4 | MR24 | M242-2 | SnTox8 | 2 dpi | Mutant |
| 5 | MR25 | M538-2 | SnTox8 | 2 dpi | Mutant |
| 6 | MR26 | M538-2 | SnTox8 | 2 dpi | Mutant |
| 7 | MR27 | M243-1 | SnTox8 | 2 dpi | Mutant |
| 8 | MR28 | M243-1 | SnTox8 | 2 dpi | Mutant |
| 9 | MR29 | M193-2 | SnTox8 | 2 dpi | Control |
| 10 | MR30 | M193-2 | SnTox8 | 2 dpi | Control |

In order to assess transcriptional differences between the mutant and wild-type lines, genes were compared and classified as upregulated, downregulated, no change, or not detected based on log_2_fold change and adjusted p-value thresholds. Transitions between these categories in the two experiments were then visualised using a Sankey plot in ggalluvial 0.12.5, providing an overview of shared and divergent regulatory responses.

### Data availability

All scripts and analyses used in this study are available in the associated GitHub repository (https://github.com/AAGI-AUS/wheat-snTox8-paper). The raw sequencing reads have been submitted to NCBI under temporary submission ID SUB16018434 and Bio Project PRJNA1432443. The address is https://www.ncbi.nlm.nih.gov/bioproject/1432443

## RESULTS

### Morphological and microscopic observations of the leaf surface after SnTox8 infiltration

The phenotypic differences between the SnTox8-sensitive wheat variety Mace and the insensitive variety Lancer following infiltration with SnTox8 and control across multiple time points are illustrated in Figure 1.A. No visible phenotypic changes were observed in either Mace or Lancer leaves up to 4 days post-infiltration under both SnTox8 and control treatments. In the sensitive line, Mace, chlorosis became visible 7 days post-infiltration (7 dpi), progressing to pronounced necrotic lesions by day 10. In contrast, Lancer showed no visible symptoms with either treatment. Similarly, Mace leaves infiltrated with control showed no phenotypic changes throughout the period. These results clearly demonstrated a differential response between Mace and Lancer to SnTox8, highlighting the distinct visible responses of the two varieties (Figure 1.A).

**Figure 1:**
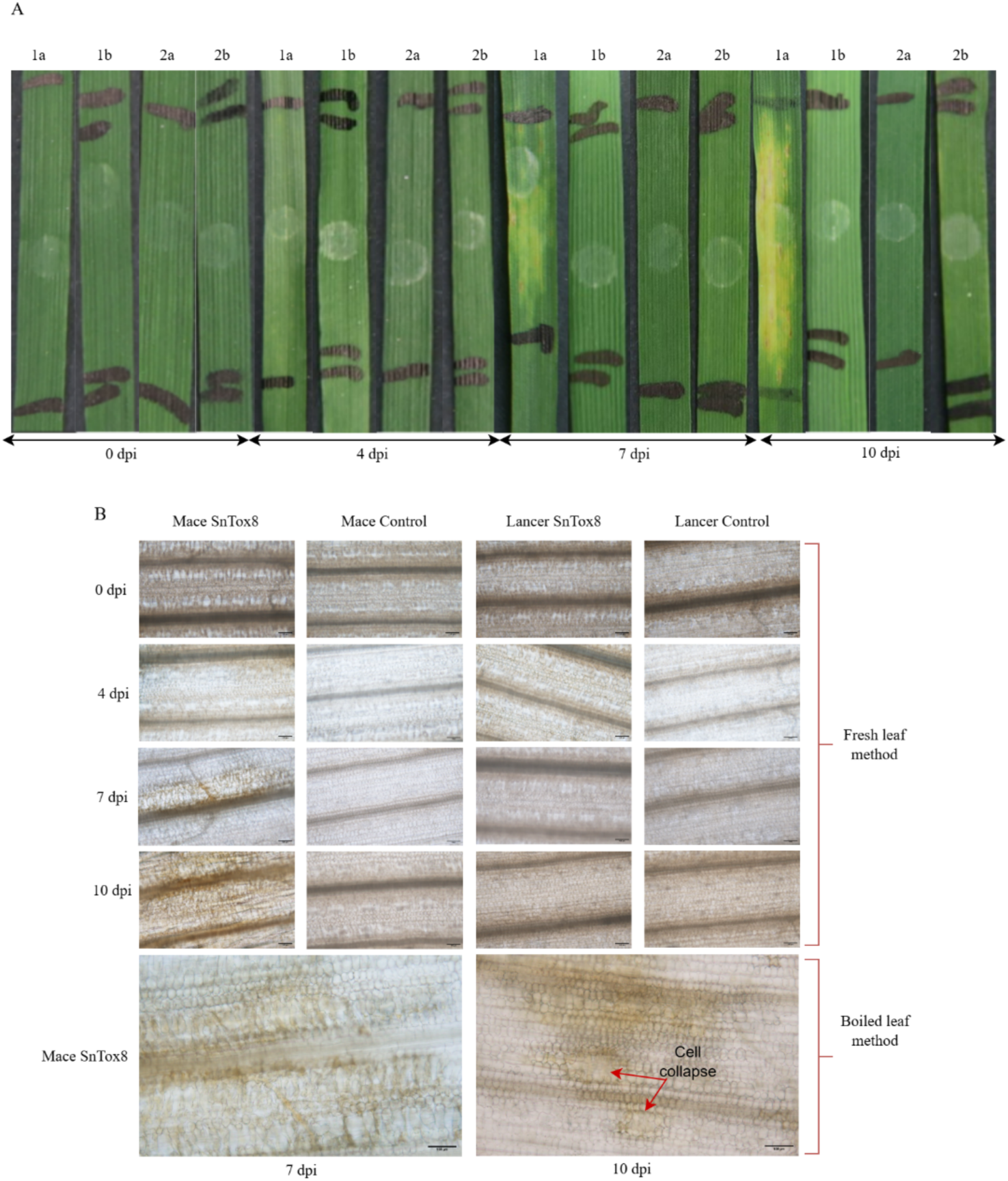
(A) Morphological observation of leaf surface of SnTox8-sensitive wheat line Mace (1) and SnTox8-insensitive wheat line Lancer (2) after SnTox8 (a) and control (b) infiltration, over time and (B) microscopic observation of Mace and Lancer leaf structure after SnTox8 and control infiltration using fresh- and boiled-leaf methods; scale bar:100µm.

Microscopic observations of symptom development over time in Mace and Lancer following infiltration with SnTox8 and the control are presented in Figure 1.B. They depict Mace and Lancer leaves immediately after infiltration, at days 4, 7, and 10 post-infiltration (dpi), using both fresh and boiled leaf preparations. No structural changes were observed in the upper surface or lower surface of the leaf in the control Mace and Lancer lines throughout the time course. Similarly, no phenotypic or structural changes were observed on the leaf surface of the SnTox8-infiltrated Lancer leaves over time. SnTox8-infiltrated Mace leaves exhibited structural changes beginning 7 dpi with visible yellowish-brown patches on the upper and lower leaf surfaces in both fresh leaf and boiled leaf treatments of SnTox8-infiltrated Mace. By 10 dpi, symptoms reached their peak in these SnTox8-infiltrated Mace samples, with necrotic lesions and cell destruction evident, and showing significant collapse of the upper-surface mesophyll cells in both treatments; despite these changes, the lower-surface stomata remained visible. These observations provide clear evidence of structural damage confined to the sensitive Mace leaves.

### Gene expression overview for RNA-seq samples

Mace and Lancer were infiltrated with either SnTox8 or control to examine transcriptional responses over five time points. Principal component analysis (PCA) on the gene counts reveals a clear separation between 0 dpi and the subsequent timepoints in the Mace (PC1), covering 80% variance. At 2 and 3 dpi, control samples are distinguishable from those treated with SnTox8, showing a clear separation at PC2 with 16% variance. This clear separation between SnTox8-treated and control samples at 2 and 3 dpi indicates that significant changes in gene expression occurred at these early time points, reflecting the plant’s initial recognition and response to SnTox8. SnTox8-treated samples continue to cluster together away from the control samples at 5 and 7 dpi. Within this cluster, the 2 and 3 dpi SnTox8-treated samples were positioned closer to the 5 and 7 dpi control samples, while the 5 and 7 dpi SnTox8-treated samples were distant from the rest (Figure S1). In Lancer, 0 dpi samples are also separate from the later time points. However, at 2, 3, 5, and 7 dpi, no clear separation between treatment and control was observed; their replicates clustered together irrespective of condition (Figure S1).

### Differential gene expression dynamics in response to SnTox8

After filtering out lowly expressed genes, a total of 67,611 genes in Mace and 66,995 genes in Lancer were retained for differential expression analysis. Figure 2 presents the Volcano plots and Upset plots of differentially expressed genes (DEGs) at each time point for both Mace (Figure 2 - A, B, C, D, and I) and Lancer (Figure 2 - E, F, G, H, and J), highlighting the upregulated and downregulated genes. In Mace, among the time points, 3 dpi showed the highest number of DEGs (8,351), followed by 2 dpi (7,387) and 7 dpi (7,168). 5 dpi has the lowest number of DEGs, 3,952. Notably, at each time point, the number of upregulated DEGs was substantially greater than the number of downregulated DEGs. At 3 dpi, the majority of DEGs were upregulated (6,502), and there were many fewer downregulated genes (1,849). A similar trend was also observed at 2, 5 and 7 dpi. In total, 12,679 unique DEGs were detected in at least one of the time points in the sensitive line, Mace. Among these, 2,052 DEGs were identified across all four time points, indicating a subset of commonly regulated genes, as shown in the *UpSet* plot in Figure 2.I. At 2 dpi and 3 dpi, a total of 1,938 genes were commonly expressed and unique only to these two timepoints, representing the second-highest overlap among all comparisons and suggesting a shared transcriptional response between these two stages of infection. In addition, 1,227, 1,420, 438, and 1,838 DEGs were unique to 2, 3, 5, and 7 dpi, respectively. Timepoints 2 dpi and 3 dpi display the largest number of unique genes shared only within these two time points, which pinpoints these two time points as an early response reaction (Figure 2. I).

**Figure 2:**
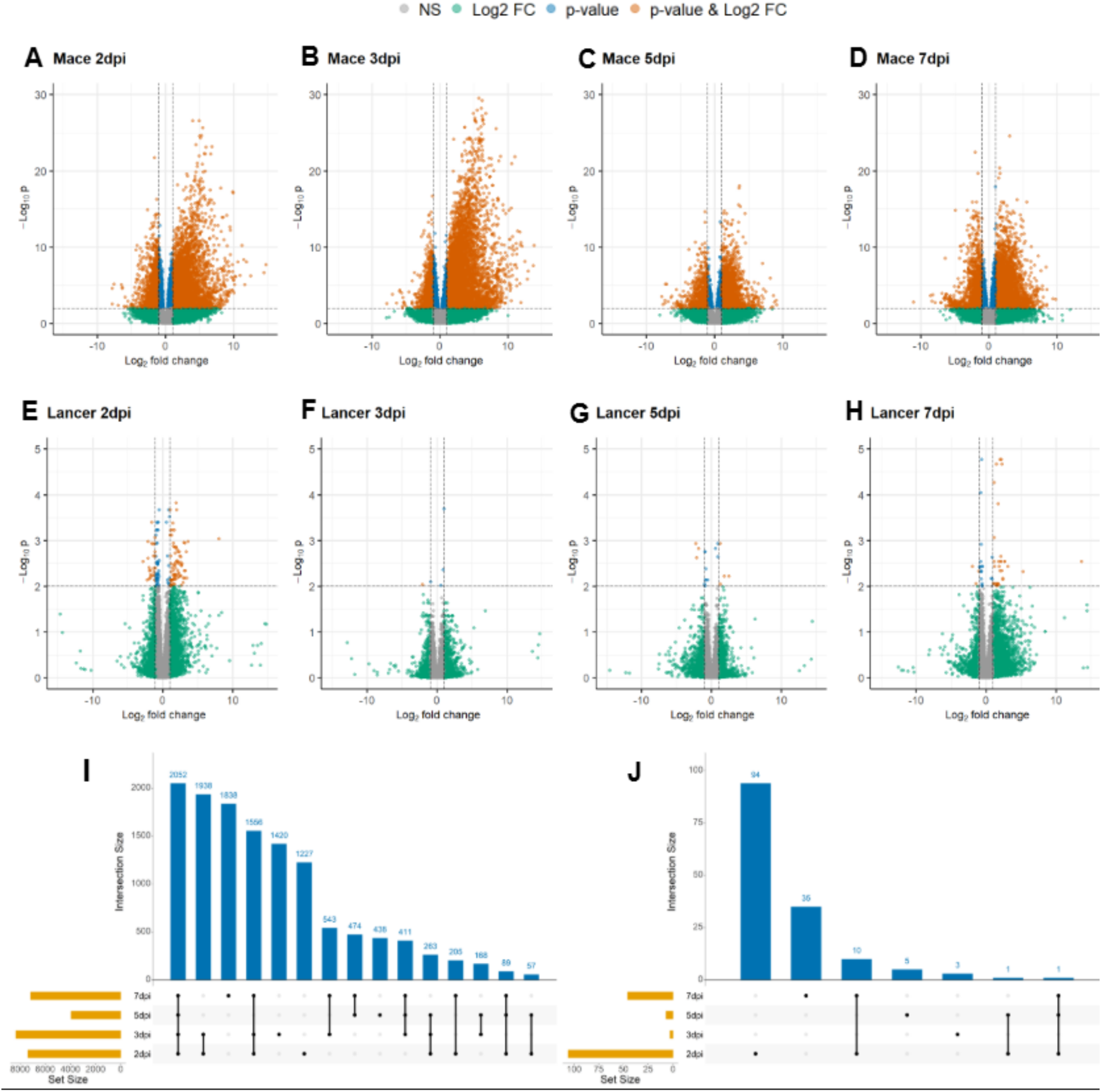
Volcano plots of differentially expressed genes (DEGs) in wheat lines Mace and Lancer at 2, 3, 5, and 7 days post SnTox8 infiltration (dpi). In Mace, a pronounced upregulation of genes is observed at 2 and 3 dpi, with a more moderate response at 5 and 7 dpi (A, B, C, D respectively), while Lancer shows comparatively limited transcriptional changes across all time points (E, F, G, H) and UpSet plots showing DEG overlap in Mace (I) and Lancer (J), with Mace displaying the largest number of unique genes at 2 and 3 dpi, highlighting a common expression pattern at these stages.

In contrast to Mace, only a small number of DEGs were identified in the insensitive line, Lancer. A total of 149 unique genes were detected across the 2, 3, 5, and 7 dpi time points, and each time point has a total of 106, 3, 7, and 46 DEGs, respectively. Among these, no DEGs were common to all time points. Instead, 94, 3, 5, and 35 DEGs were uniquely detected at 2, 3, 5, and 7 dpi, respectively (Figure 2.J). These results highlight the distinct gene expression response of Mace at both early and late time points. The minimal number of DEGs observed in Lancer again suggests a lack of recognition and response to SnTox8.

The total of 12,679 unique differentially expressed genes in Mace and 149 in Lancer were clustered based on their expression profiles (Figure S2). The resulting clusters in Mace revealed that DEGs from all samples at time point zero, including both SnTox8 and control samples, clustered together, exhibiting a similar expression at time point zero. Control samples at timepoints 2 dpi and 3 dpi formed a distinct cluster, and control samples from timepoints 5 dpi and 7 dpi clustered together, indicating that, over time, the plant recognised the control injection. In contrast, SnTox8-infiltrated samples at 2 and 3 dpi formed a distinct cluster that further grouped with the control samples at 5 and 7 dpi. Meanwhile, SnTox8-infiltrated samples at 5 and 7 dpi formed separate clusters, indicating a shift in expression over time (Figure S2.A). Unlike Mace, the control and SnTox8-infiltrated samples in Lancer displayed more similar expression profiles, suggesting a similar response between treatment and control (Figure S2.B).

Accordingly, DEGs from the sensitive line Mace and the insensitive line Lancer were used for further Gene Ontology (GO) enrichment pathway analysis.

### GO term enrichment analysis for differentially expressed genes

We performed GO enrichment analysis of DEGs at 2, 3, 5, and 7 dpi for both sensitive and insensitive cultivars. Sensitive Mace showed many significant terms across all time points, including up-regulated and down-regulated sets, which were analysed separately (Table S1, https://doi.org/10.6084/m9.figshare.32846648). Insensitive Lancer showed few or no enriched terms, indicating a minimal transcriptional response; therefore, we focused downstream analyses on Mace.

Notably, among the Mace biological process (BP) GO terms, as illustrated in Figure 3.A, enriched in the upregulated gene set, those involved in ‘transport and cellular trafficking’, ‘developmental and environmental cues’, ‘metabolism, secondary compounds’, and ‘protein and cellular organisation’ showed induction at most of the time points. Gene sets grouped into the ‘stress, defence and hormonal signalling’ were also differentially expressed, indicating the recognition of stimuli and induction of plant defence responses. Protein phosphorylation, recognition of pollen and cell surface receptor signalling pathways consistently emerged as the top-ranked categories across all time points, highlighting their central role in receptor recognition and defence-associated signalling response. We also observed that several defence-associated metabolic pathways were differentially enriched among the top-ranked terms following SnTox8 infiltration. Among them, L-phenylalanine metabolism and aromatic amino acid biosynthesis were activated, indicating enhanced flux into the phenylpropanoid pathway, which produces lignin, flavonoids, phytoalexins, and a variety of other aromatic metabolites essential for structural reinforcement and antimicrobial defence (Vogt, 2010). In addition to top-ranked terms, some pathways were consistently enriched across all time points, highlighting key components of the SnTox8-induced defence response in wheat, such as exocytosis, defence response to other organisms, and the chitin catabolic process. Significant gene counts for these terms were included, along with p-values and Rich Factors (Table S1, https://doi.org/10.6084/m9.figshare.32846648).

**Figure 3:**
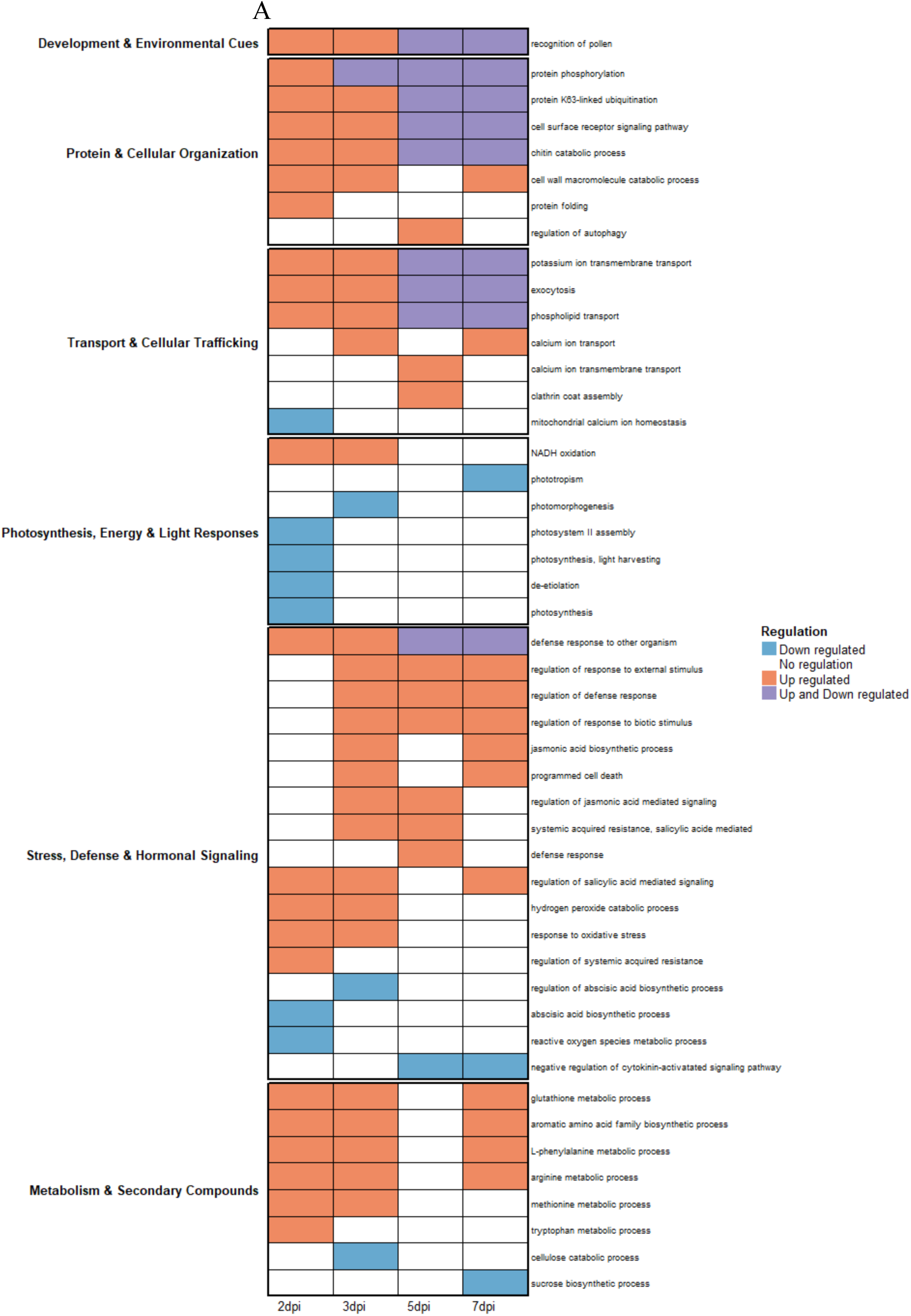
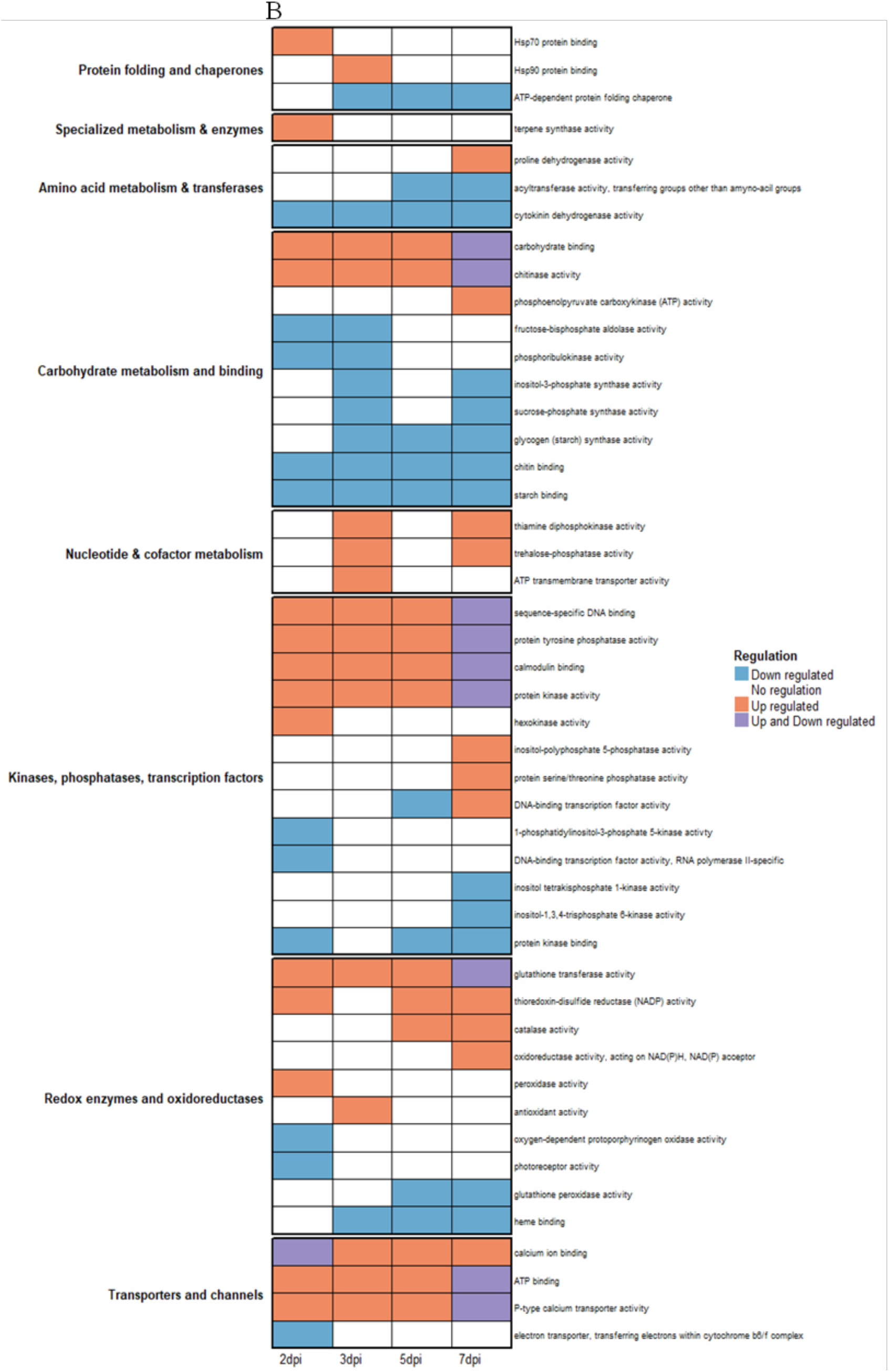
GO enrichment analysis of Mace over time. (A) Biological processes and (B) Molecular functions at 2 dpi, 3 dpi, 5 dpi and 7 dpi are involved in SnTox8 susceptibility.

Some terms were uniquely enriched in the early and later stages. Several defence-related terms, such as hydrogen peroxide catabolic process, response to oxidative stress, tryptophan metabolic process, and RO species metabolic process, were enriched only in early stages (2-3 dpi). Once recognised, the SnTox8 toxin at the early stage (2 dpi) led to downstream upregulation of defence and trafficking pathways from 3 dpi to 7 dpi, notably regulating hormone signalling (Salicylic acid, Jasmonic acid), responses to biotic/external stimuli, systemic acquired resistance, and programmed cell death were among those detected.

Conversely, GO enrichment analysis of downregulated DEGs over time revealed a marked suppression of ‘photosynthesis’ and ‘primary metabolic pathways’, indicating reduced photosynthetic activity and metabolic processes as the defence response intensified, especially at the early stage (Figure 3.A), consistent with the visible collapse of plant tissue, chlorosis and necrosis at the later stage following SnTox8 infiltration.

Consistent with the BP findings, the molecular function (MF) enrichment of upregulated genes in Mace further underscores the activation of signalling, regulatory, and defence-related biochemical activities following SnTox8 infiltration (Figure 3.B). Interestingly, gene sets are grouped into 8 categories: nucleotide and cofactor metabolism; amino acid metabolism and transferases; redox enzymes and oxidoreductases; kinases, phosphatases and transcription factors; carbohydrate metabolism and binding; protein folding and chaperones; transporters; and specialised metabolism and enzymes. Top-ranked among the enriched categories were protein kinase activity and ATP binding at all time points, reflecting strong engagement of phosphorylation-mediated signal transduction cascades that orchestrate downstream defence responses. These terms are closely aligned with the BP term, protein phosphorylation, together suggesting that phosphorylation-dependent pathways form a core component of the SnTox8-triggered defence response.

Other highly enriched functions included calcium/calmodulin binding, carbohydrate binding, and sequence-specific DNA binding. The prevalence of calcium/calmodulin binding functions supports the notion of dynamic Ca^2+^ fluxes as secondary messengers during early immune signalling (Ranty et al., 2016), while transcription factor-associated DNA binding points to transcriptional reprogramming in response to effector perception (Buscaill & Rivas, 2014). These results indicated that SnTox8 exposure triggers tightly coordinated kinase-mediated signal amplification, calcium-dependent regulation, and transcriptional control, alongside metabolic adjustments supporting defence.

In contrast, the MFs enriched among downregulated genes highlighted the suppression of energy-intensive and primary metabolic activities. At early time points (2 to 3 dpi), functions linked to carbohydrate and energy metabolism, such as fructose-bisphosphate aldolase, phosphoribulokinase, and fructose-1,6-bisphosphate phosphatase (Figure 3. B) were significantly reduced, indicating a decline in photosynthetic and carbon assimilation capacity. This early repression of photosynthetic enzymes aligns with the downregulation of photosynthesis-related pathways, as observed in the BP. At later stages (5 to 7 dpi), downregulation of MF terms extended to lipid transfer and redox-related enzymes, including acyltransferase activity and multiple inositol phosphate kinases, suggests diminished membrane trafficking and oxidative signalling as defence responses mature.

The most significantly enriched cellular components (CC) included membrane, extracellular matrix, exocyst, photosystem II oxygen-evolving complex, and chloroplast, which collectively serve as key sites for signal perception, transduction, stress-related metabolic regulation, and impact on the photosynthetic machinery during SnTox8 infiltration. Please refer to Table S1 (https://doi.org/10.6084/m9.figshare.32846648) for further details on both Mace and Lancer

### Functional characterisation of transcriptomic candidates by mutagenesis

To functionally validate the transcriptomic patterns observed in the initial RNA-Seq experiment (Experiment 1), a second RNA-Seq experiment was conducted using three M4-Mace mutant lines (Mu1, Mu2, Mu3), each with two technical replicates, infiltrated with SnTox8 (and a control) at 2 days post-infiltration (2 dpi). Here, we used both sensitive Mace and sensitive sister mutant as controls to account for background noise caused by EMS treatment. Differential expression analysis was performed using the DESeq2 package, and results were directly compared to the wild-type (WT) Mace response at the same time point from Experiment 1.

The overall transition of gene expression categories between the WT Mace and the mutants is shown in Figure 4.A, highlighting substantial shifts from upregulated to downregulated states and vice versa. As shown in the figure, WT Mace exhibited 4,699 upregulated and 2,517 downregulated DEGs following SnTox8 infiltration. In contrast, the M4-Mace mutants displayed a broader transcriptional response, with 5286 upregulated and 6536 downregulated DEGs. Comparative analysis revealed a striking divergence in gene expression between the WT and mutant lines: Of the 4,699 upregulated WT DEGs, 3,351 genes were downregulated in the mutants; of the 2,517 downregulated WT DEGs, 1,396 were upregulated in the mutants; only seven upregulated and two downregulated WT DEGs remained unchanged in the mutants. Furthermore, 1346 DEGs were upregulated, and 1114 were downregulated in WT, but showed no significant differential expression in the mutant background. These results highlight substantial transcriptional reprogramming in the mutants compared to WT Mace, supporting the notion that the mutations disrupt recognised SnTox8-triggered responses and further validating the strength of the RNA-seq findings.

**Figure 4:**
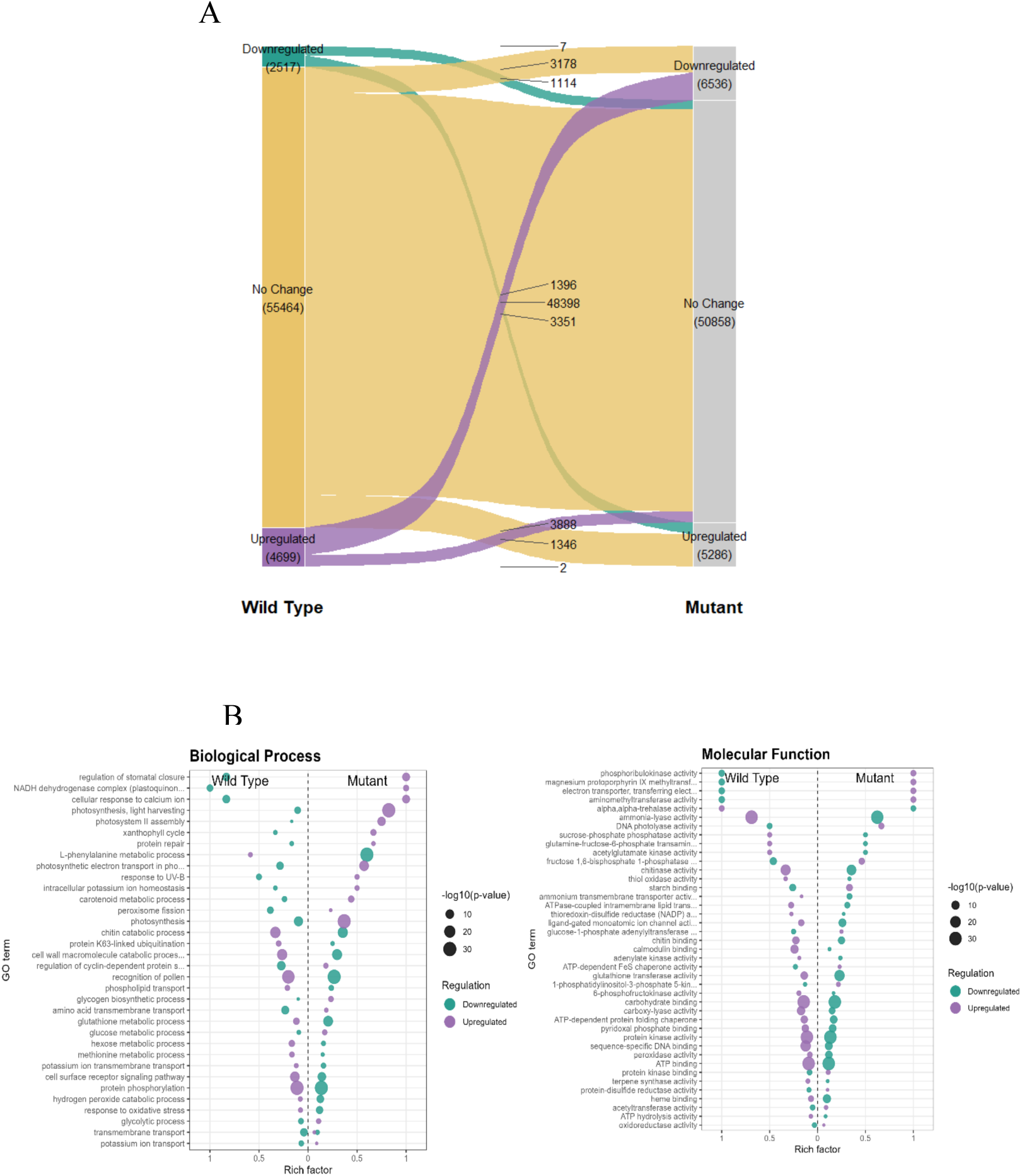
(A) Comparison between sensitive Mace-DEGs (Wild Type) and insensitive Mutant-DEGs, infiltrated with SnTox8 at 2 dpi (B) Gene Ontology enrichment analysis (BP and MF) on both the up- and down-regulated DEGs in Mace mutants.

GO enrichment analysis was conducted on both up- and down-regulated DEGs in the M4-Mace mutants and compared with WT Mace. Resulted BP and MF terms were included in Table S2 (https://doi.org/10.6084/m9.figshare.32846648). Strikingly, of the 53 significantly enriched GO (BP) terms upregulated in WT Mace, 25 were downregulated in the mutants (47.2%). Furthermore, of the 50 downregulated GO (BP) terms in WT Mace, 26 were upregulated in the mutants (52%). In addition, of the 90 significantly enriched MF GO terms upregulated in wild-type Mace, 50 (55.5%) were downregulated in the mutants, while 33 of the 66 GO terms downregulated in wild-type Mace (50%) were upregulated in the mutants. Among these, several BP and MF were identified as key enriched terms (Figure 4.B).

Among the upregulated DEGs in the mutants, the most significantly enriched biological process pathways were predominantly associated with photosynthesis and primary metabolism, suggesting upregulation of energy-generating and metabolic pathways. Conversely, the downregulated DEGs in the mutants were significantly enriched for key defence-related pathways, suggesting a possible disruption in the activation of innate immune responses to SnTox8 infiltration due to the mutated gene. Furthermore, direct defence responses that were strongly activated in sensitive Mace following SnTox8 infiltration, such as systemic acquired resistance and salicylic acid-mediated signalling regulation, were largely absent or suppressed in the mutants. Together, these results indicate that the Mace mutants fail to activate the typical SnTox8-driven defence transcriptional network, instead shifting toward metabolic programs that are inconsistent with SnTox8-triggered susceptibility, thereby validating the functional importance of the disrupted genes.

## DISCUSSION

To date, SnToxA, SnTox1, SnTox267, SnTox3, SnTox4, and SnTox5 effectors produced by *P. nodorum* have been identified. These effectors are unique to *P. nodorum* except SnToxA, which is also present in *P. tritici-repentis* and *B. polaris* (Peters Haugrud et al., 2022). Our SNB group at the Centre for Crop and Disease Management, Curtin University, has identified a new effector, SnTox8, described by Furuki et al (submitted). This study integrated transcriptomic, morphological, and microscopic analyses to investigate how two wheat genotypes, Mace (sensitive) and Lancer (insensitive), differ in their responses to the novel effector SnTox8 and in reprogramming gene expression. By integrating these datasets, we aimed to elucidate the molecular basis of SnTox8-triggered susceptibility in the sensitive genotype, Mace.

### Chlorosis and necrosis are specific to sensitive interaction

Morphological assessments of the sensitive Mace and the insensitive Lancer upon infiltration with semi-purified SnTox8 protein revealed distinct phenotypic differences between the two cultivars. In Mace, symptoms were apparent as chlorosis of the laminae at 7 dpi and progressed to necrotic lesions at 10 dpi. Lancer remained asymptomatic across all time points and treatments, confirming its insensitivity to SnTox8. The asymptomatic nature in Lancer, even at later stages, suggested that this genotype lacks the ability to recognise SnTox8, thereby conferring resistance. Microscopic examination of SnTox8-infiltrated wheat leaves further confirmed that SnTox8 specifically induces tissue damage only in the sensitive genotype. In Mace, structural alterations became apparent by 7 dpi, with the appearance of yellowish-brown lesions, consistent with chlorosis and early cell death, and by 10 dpi, widespread tissue breakdown suggested a localised effect of SnTox8. This supports the hypothesis that SnTox8 acts as a necrotrophic effector that exploits host cell death for disease development, in line with the inverse gene-for-gene model.

### Transcriptomic profiling revealed host recognition of SnTox8

The transcriptomic profiling of SnTox8-infiltrated Mace and Lancer leaves reveals contrasting molecular responses, reflecting the phenotypic sensitivity observed at the tissue level. In Mace, principal component analysis (PCA) showed clear separation between SnTox8-treated and control samples at 2 and 3 dpi (16% variance), indicating significant changes in gene expression. These early responses likely reflect initial recognition of SnTox8 and the activation of defence-like pathways, which later lead to chlorosis and necrosis in the sensitive genotype.

By contrast, Lancer showed no clear separation between SnTox8-treated and control samples at 2, 3, 5, and 7 dpi, with replicates clustering together regardless of condition. This indicates an absence of effector perception or downstream signalling, consistent with the cultivar’s insensitive phenotype.

During invasion, pathogens face sophisticated surveillance systems, called PTI and ETI, which detect and respond to potential threats before they cause serious damage. One major layer of this defence involves PRRs (pattern recognition receptors) located on the surface of plant cells. These receptors act like security sensors, scanning for PAMP molecules that are characteristic of many microbes but absent in the plant itself. These PRRs specifically detect PAMPs such as chitin, a structural component of fungal cell walls. In our study, we found that upregulation of genes related to the cell-surface receptor signalling pathway and chitin catabolic process, indicating that SnTox8 toxin detection serves as a danger signal, thereby activating defence mechanisms. Additionally, during infection, plants produce chitinases as part of their defence process, which break down fungal chitin into smaller oligomers (Dobon et al., 2016). Further, we observed pollen recognition as one of the top-ranked terms across all time points, a new finding in *P. nodorum* toxin-wheat interactions. This may reflect activation of recognition receptors that evolved from or share domains with pollen or “foreign material” recognition systems. Validation of these term-related genes should be conducted in future research to confirm whether these receptors are PRRs. Once recognised, PAMPs/MAMPs trigger the first line of defence, the PTI mechanism.

### SnTox8 triggers PTI/ETI cascades leading to ROS production and cell death

PTI and/or ETI mechanisms ultimately trigger downstream immune responses, including production of reactive oxygen species (ROS), Ca^2+^ influx, MAPK activation, synthesis of defence hormones, defence gene expression, and programmed cell death (Jones & Dangl, 2006; Peters Haugrud et al., 2019; Ramírez-Zavaleta et al., 2022). We observed such defence-related pathway upregulated, including the ROS metabolic process (3 dpi), calcium signalling pathways (all timepoints), defence response pathways (all timepoints), and programmed cell death (3 and 7dpi), confirming the hypothesis that SnTox8 effector protein triggers host cell death responses to facilitate the pathogen’s necrotrophic infection cycle (Figure 3).

The production of ROS is among the earliest described cellular responses to pathogen invasion. While early ROS production can aid resistance against both biotrophs and necrotrophs, at later infection stages it promotes resistance in both biotrophs and hemibiotrophs via regulated cell death. However, it enhances necrotrophic virulence (Ghozlan et al., 2020). ROS are important in activating defence by reinforcing cell walls, regulating defence genes, and producing toxic free radicals. Plasma membrane-located NADPH oxidase, amine oxidase, and cell wall-bound peroxidases are the potential enzyme sources for ROS (Adhikari et al., 2009; Bolwell & Wojtaszek, 1997). Comparing our results with previous findings, several ROS-related activities, such as glutathione transferase activity, peroxidase activity, Hsp90 protein binding (Adhikari et al., 2009), and proline dehydrogenase activity (Cecchini et al., 2011; Fabro et al., 2016), were found in sensitive Mace at different timepoints. Notably, the Hsp90- and Hsp70-binding activities identified in this study represent key heat shock protein functions. These molecular chaperones were reported to play pivotal roles in maintaining protein homeostasis and supporting plant survival under heat-stress conditions (Song et al., 2025). In addition, typical markers of ROS activity, such as oxidative burst, redox regulation, and detoxification mechanisms, also provide a good indication of plant’s defence against necrotrophic fungal infection (Bolwell & Wojtaszek, 1997).

Evidence of ROS production was clear in our study. Its activities; as indicated in other studies such as upregulation of antioxidant activity (Foyer & Hanke, 2022), hydrogen peroxide catabolic process (Adhikari et al., 2009), downregulation of glutathione peroxidase activity, upregulation of catalase, thioredoxin-disulphide reductase (NADP), and oxidoreductase activity (Bolwell & Wojtaszek, 1997; Foyer & Hanke, 2022), and calcium ion binding was found at any time point in our study (Figure 3), leading to cell death. In a similar study, ROS served as key signalling molecules that trigger cell death in ToxA-sensitive wheat, contributing to necrosis during infection by *Pyrenophora tritici-repentis* (Adhikari et al., 2009), consistent with our findings.

### Regulation of effector-induced cell death via protein phosphorylation pathways

The death of an invaded cell, known as the hypersensitive response, is usually characterised by rapid, transient responses occurring mainly at the plant cell surface and based predominantly on the activation of pre-existing components, such as changes in protein phosphorylation patterns, and ion fluxes (Bolwell & Wojtaszek, 1997). Protein phosphorylation regulates protein functions by transferring the phosphate from ATP to the hydroxyl residue of the protein substrate with the help of a protein kinase (Hanks et al., 1988; Kesawat et al., 2022; Shaw et al., 2014), while a protein phosphatase facilitates the reverse reaction (Phan et al., 2021). Research has showed that protein phosphorylation cascades and calcium fluxes are involved in necrotrophic fungal effector-induced cell death (Rasmussen et al., 2004). In the present study, protein phosphorylation, along with protein kinase activity, was identified as the top pathway at all timepoints in sensitive wheat. Evidence includes many phosphatases and kinase activities, upregulated activities included protein tyrosine phosphatase (all timepoints), trehalose-phosphatase (3 and 7 dpi), thiamine diphosphokinase (3 and 7 dpi), protein serine/threonine phosphatase (7 dpi), inositol-polyphosphate 5-phosphatase (7 dpi), and phosphoenolpyruvate carboxykinase (ATP) (7 dpi). These phosphatases are known to fine-tune signal transduction by dephosphorylating key signalling proteins, thereby modulating MAPK cascades, hormone signalling, and stress responses (Luan, 2003; Shi, 2009). For example, trehalose-phosphatase activity regulates trehalose-6-phosphate levels, which act as a sugar-signalling molecule linking metabolism with development in response to carbon availability and stress adaptation (Paul et al., 2008).

### Suppression of phosphoinositide signalling at the late stage of infection

Inositol phosphate kinases (IPKs) are enzymes that regulate the production of inositol polyphosphates. These important signalling molecules act as second messengers in eukaryotic cells. Because these molecules control many essential cellular processes, understanding the genes that produce them is biologically important. However, the IPK gene family has not been well studied in bread wheat (*Triticum aestivum*). Gene ontology analysis showed that wheat IPKs are involved not only in inositol phosphate signalling, but also in various metabolic and biological processes, suggesting broad functional roles. Under normal conditions, inositol phospholipids (phosphoinositides, PIs) play a crucial role in activating plant immunity. Their activities are induced when plants detect a pathogen via immune receptors such as FLS2, PIs. This triggers downstream defence responses, including calcium release, ROS generation, and the hypersensitive response (HR). Pathogens are known to manipulate this inositol phosphate metabolic pathway to evade NB-LRR recognition and enhance the virulence function of effectors (Kale et al., 2010). One example of that is *Phytophthora infestans* effector Avr3a which binds to lipid phosphatidylinositol phospholipase C substrates PI3P, PI4P, and PI5P, stabilising itself within plant cells and enhancing virulence (Yaeno et al., 2011). However, several enzymes in the inositol kinase family in our study were downregulated at the late timepoint (7 dpi), including inositol tetrakisphosphate 1-kinase, inositol-1,3,4-trisphosphate 6-kinase, inositol-1,3,4-trisphosphate 5-kinase, and 1-phosphatidylinositol-4-phosphate 5-kinase (Table S1, https://doi.org/10.6084/m9.figshare.32846648). Suppression of these kinases could impair calcium-dependent defence signalling and vesicle trafficking, potentially affecting necrotrophic colonisation (Hung et al., 2014). Similar results were also found in other pathosystems; transcriptomic analyses of powdery mildew and rust pathogens (obligate fungal biotrophs) in wheat revealed that suppression of the phosphoinositol signalling pathway occurs in susceptible lines (Poretti et al., 2021).

Overall, the late-stage suppression of key inositol kinase genes in SnTox8-sensitive wheat suggests that the disruption of phosphoinositide signalling may reflect a host-driven response that ultimately weakens defence activation and limits HR-associated immunity. This reduction in signalling may also represent a metabolic adjustment, whereby the plant down-regulates energy-intensive pathways during later infection stages to balance the substantial energetic costs of maintaining prolonged immune responses.

### SnTox8-mediated suppression of photosynthesis and primary metabolism

Signalling pathways such as photosynthesis, photosystem II assembly, photosynthetic electron transport, and photomorphogenesis were downregulated at early time points (2 and 3 dpi), and the sucrose biosynthetic process was downregulated at later time points (7 dpi), indicating that SnTox8 protein suppresses or degrades photosynthetic machinery. Similar results were found from transcriptomic studies of wheat responses to the necrotrophic effector SnToxA, an interaction found in both *P. nodorum-* and *Pyrenophora tritici-repentis*-wheat pathosystems, as well as for SnTox3 from *P. nodorum*. SnToxA primarily disrupts the Calvin cycle, while SnTox3 interferes with photosystem I and ATP synthase, potentially leading to feedback inhibition of photosynthetic activity (Winterberg et al., 2014). Also, it was evident that photosystem II is a major source of the secondary oxidative burst in HR-related resistance and observed notable change of downregulation of several chloroplast-associated genes in response to necrotrophic effector ToxA, stating the inhibition of photosystem activity in the chloroplast by this effector; thus leading to high levels of ROS that can impact cellular homeostasis (Adhikari et al., 2009; Manning et al., 2009). This supports the hypothesis that the chloroplast photosystem of toxin-sensitive plants serves as a primary source of ROS in this interaction.

### SnTox8 exploits hormonal signalling to promote necrotrophic infection

Since plant R-genes are not associated with resistance to necrotrophs, defence relies on the balanced regulation of phytohormones, jasmonic acid (JA), ethylene, salicylic acid (SA), and abscisic acid (ABA) (Birkenbihl & Somssich, 2011; Ghozlan et al., 2020). Furthermore, ROS such as superoxide anion and hydrogen peroxide can trigger signalling pathways including SA synthesis and systemic acquired resistance (SAR) (Ghozlan et al., 2020), a long-lasting, broad-spectrum defence mechanism associated with SA accumulation (Ezzat et al., 2014). Hormonal signalling pathways, such as systemic acquired resistance, salicylic acid, and jasmonic acid-mediated signalling pathways (2, 3, 5 dpi), are upregulated in early timepoints in our study. According to Ghozlan et al (2020), elevated SA enhances resistance to hemibiotrophs but increases susceptibility to necrotrophs. Conversely, JA signalling generally promotes resistance and systemic immunity against necrotrophs. ABA’s role is more variable, acting as either a positive or negative regulator depending on the specific host-pathogen interaction (Ghozlan et al., 2020). Our study shows downregulation of the ABA biosynthetic pathway at 3 dpi.

### SnTox8-induced molecular pathways were confirmed by loss-of-function mutants

The extensive reversal of SnTox8-responsive gene expression in the Mace mutants indicates that the mutated loci involved in the transcriptional program associated with SnTox8-mediated susceptibility. Suppression of defence-related pathways, such as systemic acquired resistance and salicylic acid signalling, together with upregulation of photosynthesis and primary metabolic processes, suggested that the mutants fail to recognise the SnTox8 signal. These results provide functional validation that the disrupted genes (corresponding host gene to SnTox8) are required for activating host responses exploited by SnTox8 during NE infection. Similar mutagenesis-based functional validations have previously been instrumental in confirming susceptibility loci in other *P. nodorum*-wheat interactions. For instance, the cloning of *Snn1* and *Tsn1* was supported by ethyl methanesulfonate (EMS) mutant analyses, which showed loss of effector sensitivity when the corresponding receptor-like kinase or NLR-like genes were disrupted (Shi et al., 2016; Faris et al., 2010; Phan et al., 2021). Likewise, *Snn3-D1* was identified through mapping and validated by mutant screening that revealed insensitivity to SnTox3 (Zhang et al., 2021).

### Comparative transcriptomic insights into SnTox8 signalling

Comparable host analysis studies have been previously published for ToxA from both *P. nodorum* and *P. tritici repentis* (Pandelova et al., 2009) and for SnTox3 from *P. nodorum* (Winterberg et al., 2014). Both ToxA and SnTox3 studies used the GeneChip Wheat Genome Array, whereas this SnTox8-treated study used total RNA-seq analysis. Our data, consistent with the SnToxA and SnTox3 studies, indicate massive transcriptional reprogramming in response to SnTox8 treatments, including cellular responses associated with defence. This and SnToxA studies support the notion that toxin-induced cell death is triggered by impairment of the photosynthetic machinery and by the accumulation of ROS. SnTox3 also triggers the collapse of photosynthesis in a manner distinct from SnToxA as described above. In addition, both SnTox3 transcriptomic and proteomic data showed upregulation of methionine metabolism which is similar to SnTox8. SnTox8 infiltration upregulated the methionine metabolic process at the early stage of infection (2 and 3 dpi) and methionine adenosyltransferase activity at 2, 3,3 and 7 dpi. When methionine metabolism is upregulated, the plant produces higher levels of homocysteine and S-adenosyl methionine (SAM). These metabolites feed into multiple defence and cell-death-related pathways. For instance, SAM is a precursor of ethylene, a defence hormone, and is the main methyl donor for the production of many defence compounds, particularly phenylpropanoids, such as feruloylquinic acid and chlorogenic acid (Moffatt & Weretilnyk, 2001; Winterberg et al., 2014). Methionine adenosyltransferase (MAT) is the enzyme that converts methionine into SAM (Brosnan & Brosnan, 2006; Sauter et al., 2013).

We also observed significant differences in plant responses to SnTox8, SnTox3, and SnToxA, which are attributable to their effects on secondary metabolism. SnToxA strongly activated the tryptophan biosynthetic pathway, as demonstrated by microarray analyses. In contrast, SnTox3 did not induce the accumulation of tryptophan-related metabolites, consistent with microarray data showing minimal activation of genes within this pathway. SnTox8 also activated the tryptophan metabolic process as a highly enriched term in the early timepoint (2 dpi). Further, SnTox3 induced the phenylpropanoid pathway, evidenced by increased expression of associated genes and the detection of key phenylpropanoid-derived metabolites, including feruloylquinic acid and chlorogenic acid. The phenylpropanoid pathway is a major metabolic pathway in plants, producing many important defence-related secondary metabolites, including lignin, salicylates, coumarins, flavonoids, and phytoalexins. These compounds help strengthen cell walls, detoxify pathogens, or directly inhibit pathogen growth. The initial steps of the pathway for the production of these various compounds rely on the conversion of phenylalanine to precursor molecules by enzymes including phenylalanine ammonia-lyase, cinnamate4-hydroxylase, and 4-coumarate: coenzyme A ligase, all of which show a strong increase in gene expression in leaves treated with the necrotrophic effector ToxA at around 9 hours post infection, but the synthesis of flavonoid compounds and phytolalexins is unclear. SnTox8 also triggers the L-phenylalanine metabolic and catabolic processes as highly enriched top terms at 2, 3, and 7 dpi and the flavonoid biosynthetic process at 3 dpi.

Comparative transcriptomic analyses of SNB-host interactions demonstrate that although SnToxA, SnTox3, and SnTox8 all induce host cell death in susceptible wheat lines, each effector also triggers additional independent host responses, suggesting that they possess distinct functional properties. While SnTox8 shares several core defence-associated transcriptional signatures with SnToxA and SnTox3, it uniquely activates diverse metabolic pathways, particularly in secondary metabolism, setting it apart from previously characterised necrotrophic effectors. These distinctive molecular patterns indicate that SnTox8 engages a more complex network of defence signalling, metabolic rewiring, and stress-associated processes.

In conclusion, the SnTox8-wheat interaction in Mace showed strong susceptibility marked by ROS bursts, calcium signalling, phosphorylation cascades, hormonal shifts, and suppressed photosynthesis, collectively hijacking defence responses to promote cell death. The insensitive line, Lancer, exhibited minimal or no transcriptional response, suggesting no effector recognition. These contrasting profiles identify key signalling and metabolic pathways for future functional studies and resistance breeding. Looking forward, fine mapping and functional validation of the candidate susceptibility genes identified in this study, including those linked to SnTox8 perception, signalling, and metabolic responses, will be essential for elucidating the genetic basis of SnTox8-mediated susceptibility. Future work should also explore the biochemical role of SnTox8 within host tissues, dissect the crosstalk between metabolic pathways and cell-death signalling, and clarify how phosphoinositide suppression contributes to effector success. The molecular markers derived from this research will support marker-assisted selection and offer promising avenues for breeding SNB-resistant wheat cultivars. Ultimately, integrating genomic, transcriptomic, and mutational approaches will accelerate the cloning of *Snn8* and provide a more complete understanding of effector-triggered susceptibility in wheat.

## Acknowledgements

This project was supported by a co-investment between Curtin University and the Grains Research and Development Corporation (CUR00023); the data analysis was supported by the Analytics for the Australian Grain Industry (AAGI) GRDC project CUR2210-005OPX, and the PhD scholarship was funded by the Australian Research Training Program (RTP). The authors also acknowledge the Curtin Health Innovation Research Institute for providing laboratory space and technology platforms for widefield microscopy, with assistance from Mr Michael Nesbit.

## Authors’ contributions

AAPV led the conception and writing of the manuscript. KG, HP, and MG contributed to critical revision and editing of the manuscript. AAPV, KG and HP were also involved in the data analysis, while AAPV and KG also contributed to data visualisation. Assistance with SnTox8 preparation was provided by EF, and other laboratory assistance was provided by FK and KC. The EMS-Mutagenesis protocol was provided by SP. All authors read and approved the final manuscript.

## Ethics approval statement

This study did not involve human participants or animals; therefore, ethics approval was not required.

## Permission to reproduce material from other sources

All third-party material included in this manuscript has been reproduced with appropriate permission and is duly acknowledged.

## List of abbreviations

ABA: Abscisic acid
AGRF: Australian Genome Research Facility
AAGI: Analytics for the Australian Grain Industry
AGT: Australian Grain Technologies
ATP: Adenosine triphosphate
BAM Binary: Alignment/Map
BP: Biological Process (Gene Ontology)
CC: Cellular Component (Gene Ontology)
CDPKs: Calcium-dependent protein kinases
DEG(s): Differentially expressed gene(s)
DNA: Deoxyribonucleic acid
dpi: Days post-infiltration
EMS: Ethyl methane sulfonate
ETI: Effector-triggered immunity
ETS: Effector-triggered susceptibility
FLS2: Flagellin-Sensing 2
GO: Gene Ontology
GRDC: Grains Research and Development Corporation
H_2_O_2_: Hydrogen peroxide
HR: Hypersensitive response
IGV: Integrative Genomics Viewer
IPK(s): Inositol phosphate kinase(s)
JA: Jasmonic acid
KOH: Potassium hydroxide
MAPK: Mitogen-activated protein kinase
MAT: Methionine adenosyltransferase
MF: Molecular Function (Gene Ontology)
mRNA: Messenger RNA
NE(s): Necrotrophic effector(s)
NADPH: Nicotinamide adenine dinucleotide phosphate
NB-LRR(NLR): Nucleotide-binding Leucine-rich repeat
PAMP(s): Pathogen-associated molecular pattern(s)
PBS: Phosphate-buffered saline
PCA: Principal component analysis
PI(s): Phosphoinositide(s)
PRR(s): Pattern recognition receptor(s)
PTI: PAMP-triggered immunity
QTL(s): Quantitative trait loci
RNA: Ribonucleic acid
RNA-seq: RNA sequencing
ROS: Reactive oxygen species
RTP: Research Training Program
SA: Salicylic acid
SAM: S-adenosyl methionine
SAR: Systemic acquired resistance
SCN: Soybean cyst nematode
SNB: Septoria nodorum blotch
SnTox: Stagonospora nodorum toxin (effector)
Snn: Sensitivity gene in wheat
WT: Wild type

## Supporting information legends

**Figure S1:**
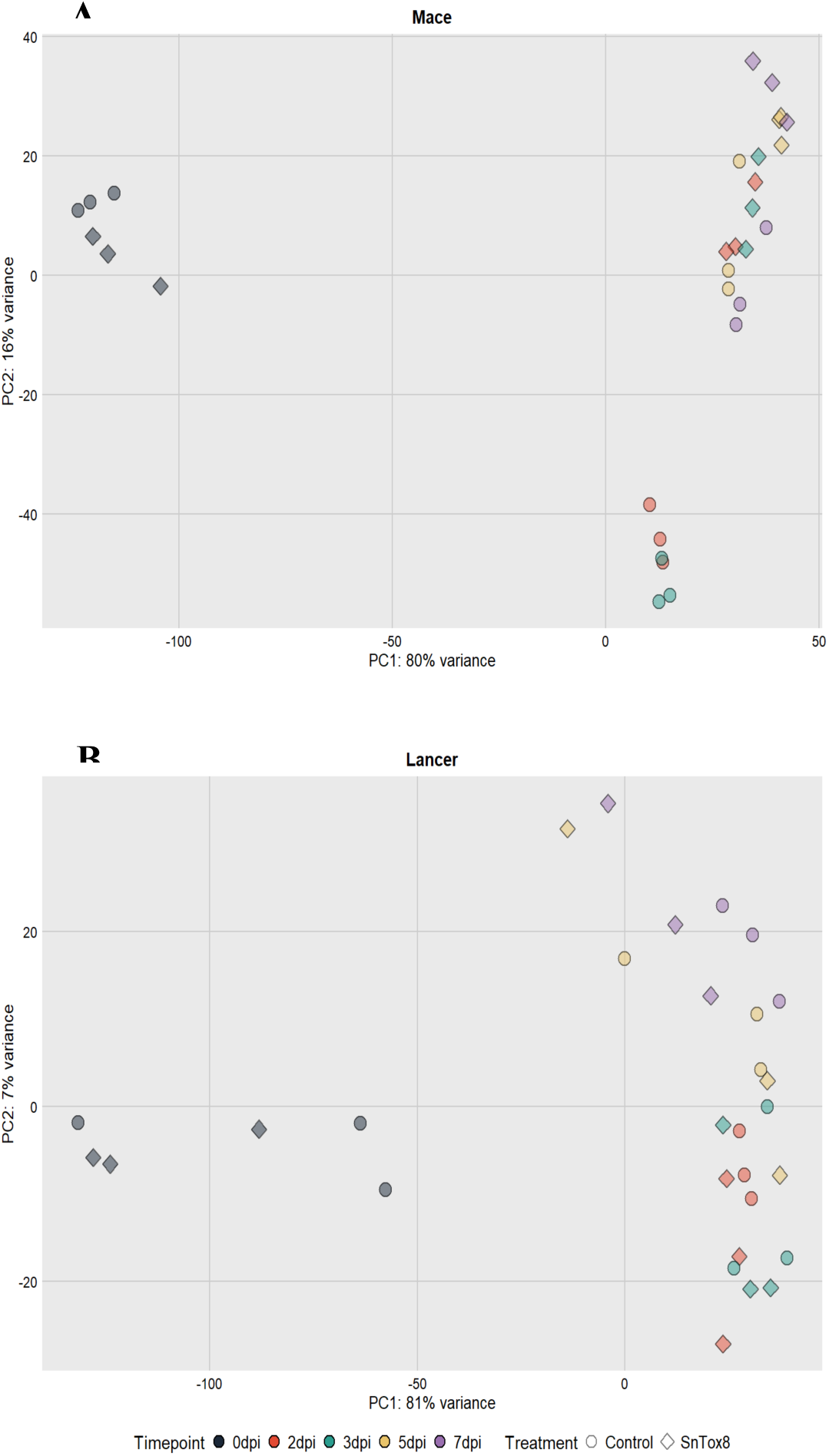
Principal Component Analysis (PCA) of gene expression in wheat lines *Mace* (A) and *Lancer* (B). In *Mace* (A), samples show clear clustering by treatment, particularly at 5 and 7 days post-infiltration (dpi), whereas *Lancer* exhibits less distinct separation (B).

**Figure S2:**
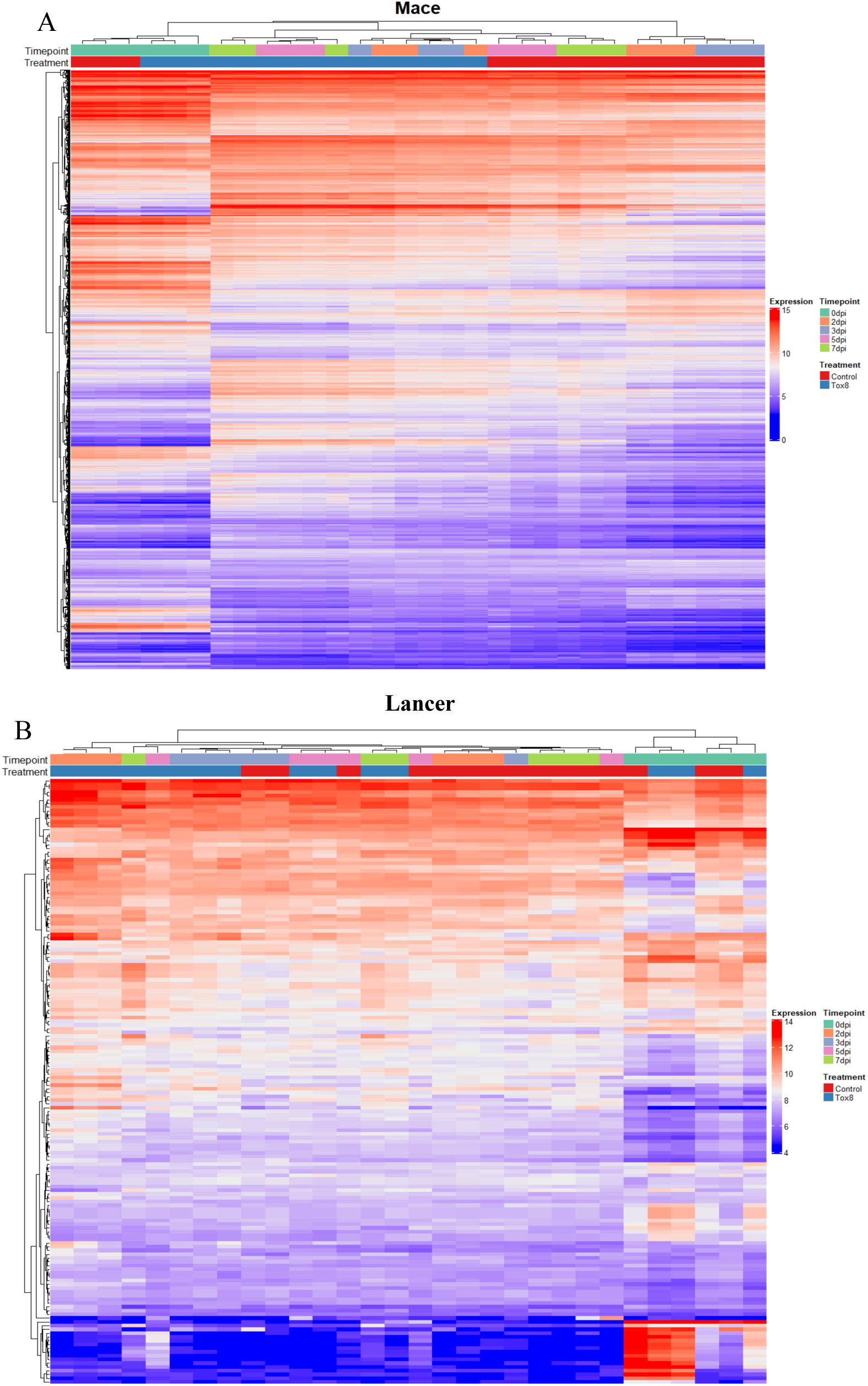
Heatmaps of DEGs in Mace (A) and Lancer (B). Mace shows clear clustering by time and treatment, while Lancer exhibits similar expression patterns across control and SnTox8-infiltrated samples.

Table S1: GO term enrichment analysis results (biological process, molecular function and cellular component) of both upregulated and downregulated DEGs of Mace and Lancer at 2, 3, 5, and 7 dpi.

Table S2: GO enrichment analysis results (biological process and molecular function) of both up- and down-regulated DEGs in the Mace mutants compared with WT Mace.

Figures and supporting information link https://doi.org/10.6084/m9.figshare.32846648

